# Midfrontal theta tACS facilitates motor coordination in dyadic human-avatar motor interactions

**DOI:** 10.1101/2021.05.25.445554

**Authors:** Sarah Boukarras, Duru Gun Özkan, Vanessa Era, Quentin Moreau, Gaetano Tieri, Matteo Candidi

## Abstract

Synchronous interpersonal motor interactions require moment-to-moment predictions and proactive monitoring of the partner’s actions. Neurophysiologically, this is highlighted by an enhancement of midfrontal theta activity. In the present study we explored the causal role of midfrontal theta for interpersonal motor interactions employing transcranial alternating current stimulation (tACS). We implemented a realistic human-avatar interaction task in immersive virtual reality (IVR) where participants controlled a virtual arm and hand to press a button synchronously with a virtual partner. Participants completed the task while receiving theta (Experiment 1) or beta (control frequency, Experiment 2) EEG-informed tACS over the frontal midline, as well as sham stimulation as a control. Results showed that frontal theta tACS significantly improved behavioural performance (by reducing interpersonal asynchrony) and participants’ motor strategies (by increasing movement times and reducing reaction times), while beta tACS had no effect on these measures. These results suggest that theta tACS over frontal areas facilitates action monitoring and motor abilities supporting interpersonal interactions.

## Introduction

The ability to coordinate our movements with those of our conspecifics is a cornerstone of successful social interactions (Sebanz et al., 2006; Vesper et al., 2010; Sacheli et al., 2012; 2015; Era et al. 2019, 2020; Gandolfo et al., 2019; Fini et al., 2020; Boukarras et al., 2021). This process relies on the anticipation of observed movements and their consequences, through motor simulation processes (Fadiga et al. 1995; Umiltà et al., 2001; Urgesi et al., 2006, 2010; Aglioti et al., 2008; Abreu et al., 2012; Avenanti et al., 2013; Candidi et al., 2014; Panasiti et al., 2017) as well as on the integration of observed and executed actions into a sensorimotor “common coding” (Jeannerod, 2000; Sebanz and Knoblich, 2009; Sacheli et al., 2015). Indeed, previous studies indicate that the simulation of observed actions occurring in the primary motor cortex is crucial for effective temporal coordination, as shown in duet piano players (Novembre et al., 2014). However, especially in conditions of high uncertainty, predictions regarding the unfolding of the partner’s actions might turn out to be incorrect and thus, need to be updated (Jacquet et al., 2016) for appropriate actions to be selected (Bekkering et al., 2009). This ability likely relies on continuous monitoring of one’s own and one’s partner’s actions.

Performance monitoring during the execution of goal-directed actions is supported by the activity of the medial frontal cortex (MFC), which signals the occurrence of errors and mismatches between expected and observed outcomes and promotes the implementation of remedial actions (Ullsperger et al., 2014). Crucially, observing someone else performing an error induces similar behavioural (Ceccarini & Castiello, 2018; Sacheli et al., 2021) and neural (Miltner et al., 2004; van Schie et al., 2004; Moreau et al., 2021) responses to those generated when performing an error in first person. EEG studies have shown that performing errors generate fronto-parietal time, i.e., Error Related Negativity (ERN) and Positivity error (Pe), and frontal time-frequency, i.e., theta Event Related Synchronization (ERS), responses (Falkenstein et al., 1991; Gehring et al., 1993; Falkenstein et al., 2000; van Schie et al., 2004; Miltner et al., 2004; Koelewijn et al., 2008; de Brujin et al., 2007; Pavone et al., 2016; Spinelli et al., 2017; Pezzetta et al., 2018). This evidence therefore makes it relevant to study the contribution of the monitoring system during interpersonal motor interaction when both self- and other-behaviour needs to be monitored and integrated in real time. A growing body of evidence indeed indicates that the MFC is involved in interpersonal performance monitoring (Bekkering et al., 2009; see Ninomiya et al., 2018 for review). Additionally, recent studies in monkeys highlighted a role of cortico-cortical premotor and medial prefrontal network for the execution of interactive tasks (Sliwa et al., 2017; Ninomiya et al., 2020). However, whether the ability to monitor and react to the behaviour of a partner causally depends on midfrontal theta remains unknown.

One recent EEG study (Moreau et al., 2020) on healthy individuals reported an increase of midline frontal theta/alpha power not only in response to observed errors (see also Moreau et al., 2021, Supplementary Materials) but also when participants need to predict and monitor the actions of a virtual partner (VP) during an interaction. Previous theories on midfrontal theta activity posit that these brain oscillations act as a nonspecific “alarm” signal that may be used to implement behavioural adjustment by synchronizing the simultaneous employment of frontal (Hanslmayr et al., 2008), motor (Nigbur et al., 2012) and sensory (van Driel et al., 2012; Fusco et al., 2020, Moreau et al., 2020) areas. Indeed, a theta phase synchrony between the MFC and other frontal sites has been repeatedly observed in various tasks eliciting the need for cognitive control (Cavanagh & Shackman, 2015).

Beside their role in conflict/error resolution, midfrontal theta oscillations have been also related to the implementation of a proactive form of cognitive control. Specifically, an increase of frontoparietal theta activity has been observed when participants are preparing to switch task compared to when preparing to repeat the same task (Cooper et al., 2015; 2017), as well as when individuals plan to avoid obstacles appearing at varying times along their walking path (Mustile et al., 2021). Moreover, trial-by-trial changes in theta power are associated with faster reaction times in switch trials, suggesting that flexible theta activity supports anticipatory cognitive control, associated to better performance (Cooper et al., 2019). Similarly, two recent studies reported the presence of pre-trial theta activity in the ventromedial prefrontal cortex during a Go/NoGo task (Adelhöfer et al., 2020) and in the anterior pre-supplementary motor area during a Stop signal task (Chang et al., 2017) associated to motor inhibition. The fact that midfrontal theta activity also supports proactive cognitive control is in line with the findings discussed above (Moreau et al., 2020), where an increase in midfrontal theta power was observed not only when participants observed a motor change in the virtual partner, but also when they predicted this change might occur during an interpersonal interaction.

To study the causal role of midfrontal theta in the ability to monitor and react to the behaviour of a partner, we designed an interactive motor task in immersive virtual reality (IVR), where participants could control a virtual body, observed from a first-person perspective, and were asked to synchronize their virtual arm and hand movements with those of a VP while receiving frontal EEG-informed transcranial alternating current stimulation (tACS, see Fig. 1). Our main goal was to test whether boosting midfrontal theta activity facilitates monitoring processes and strategic implementation of motor adjustments during interpersonal motor interactions. tACS is a non-invasive neuromodulation technique based on a weak, alternating flow of electrical current that proved to be a viable tool to entrain endogenous rhythmic activity in a frequency-dependent manner (Helfrich et al., 2014; Neuling et al., 2015; 2017) and modulate behaviour (Feurra et al., 2013; Vosskuhl et al., 2015; Fusco et al. 2018; 2020; Novembre et al., 2017). We delivered EEG-informed individualized tACS (i.e., based on participants’ individual frequencies extracted from resting-state EEG, see STAR Methods) in the theta (Experiment 1) or beta (Experiment 2) range, and a within-subject control sham stimulation over frontal midline (see Fig. 2, left panel). During tACS stimulation (approximately 9 minutes per block), two electrodes were placed over fronto-central (FCz) and parietal (Pz) positions according to the International 10-10 EEG layout (see STAR Methods). We modelled the electric field strength for our montage using the “Realistic vOlumetric-Approach to Simulate Transcranial Electric Stimulation” (ROAST), (Huang et al., 2019), see Supplementary Fig. S1. The IVR Motor Interaction Task (Fig. 2) comprised two blocks (Interactive and Cued) that differed for (i) the instruction given to the participant and (ii) the type of interaction required. Specifically, in the *Interactive* block (see Video S1), participants were asked to reach and press one of the two virtual buttons in front of them as synchronous as possible with the VP while performing either an imitative (instruction “same”) or a complementary (instruction “opposite”) movement with respect to the virtual partner’s one. If the VP performed a correction (see STAR Methods and Fig. 2), the participant needed to adapt to it by changing his/her own movement. In the *Cued* block (see Video S1), conversely, participants had to synchronize their reach-to-press movements with those of the VP as in the *Interactive* block but were in this case instructed to press either the “purple” (with the index finger) or “yellow” (with the middle finger) button, regardless of which action the avatar was performing. Thus, in the *Cued* condition, no adaptation was required in correction trials (see STAR Methods).

**Fig. 1.**
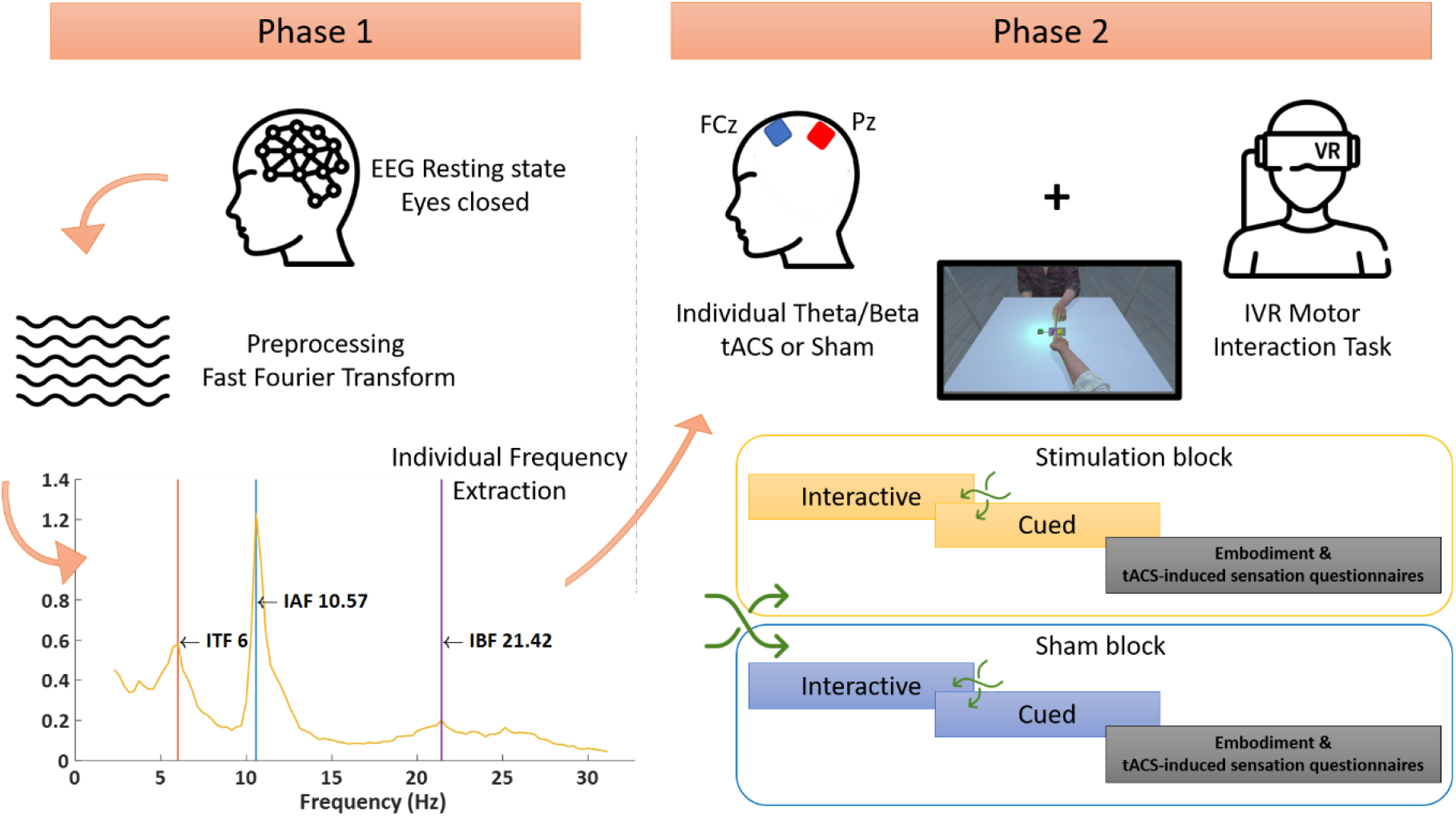
Experimental procedure. **Phase 1**: participants arrived in the laboratory and their resting-state EEG (eyes closed) was recorded for 5 minutes. EEG data were then analysed to extract the individual theta (Experiment 1) or beta (Experiment 2) frequency. **Phase 2**: participants received individualized theta (Experiment 1) or beta (Experiment 2) tACS while engaged in the IVR motor interaction task. Each stimulation block (real and sham) included one Interactive and one Cued task. The order of stimulation and task block was counterbalanced. At the end of each stimulation block, participants completed the Embodiment and tACS-induced sensations questionnaires (see STAR Methods).

**Fig. 2.**
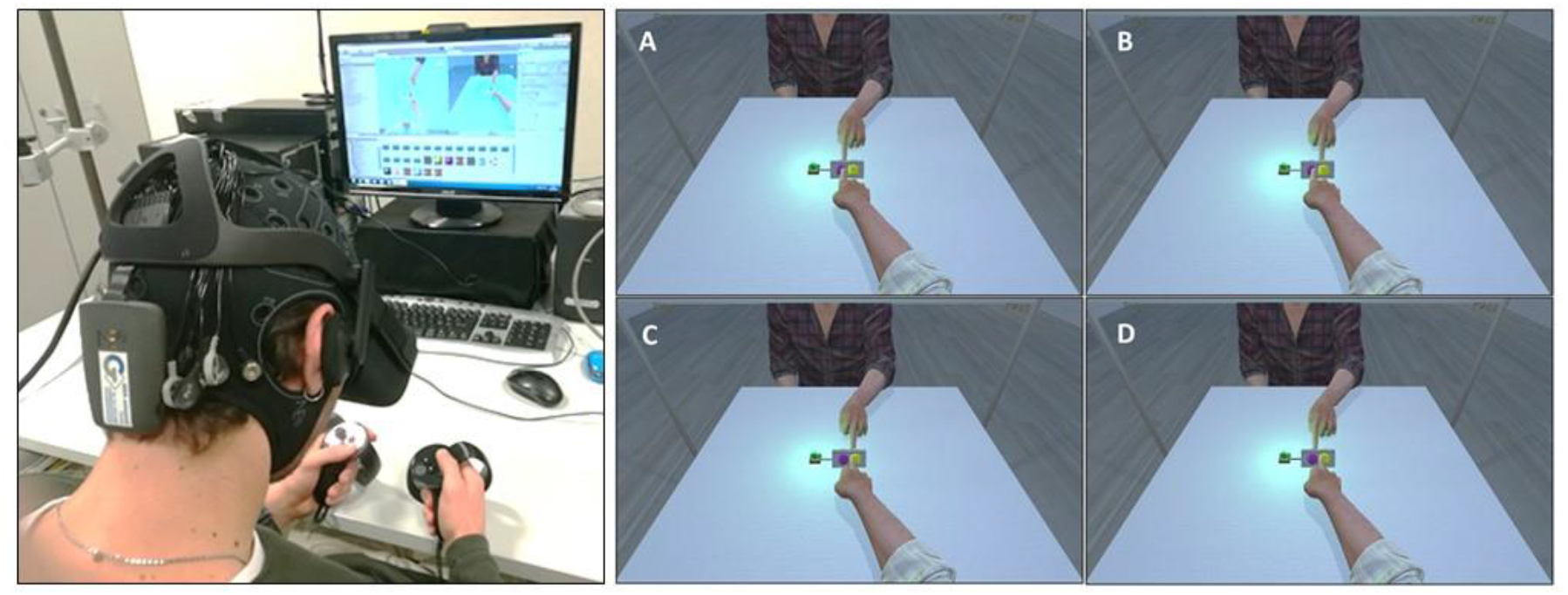
**Left Panel**: Example of the experimental setup with IVR and tACS. **Right Panel**: Motor Interaction Task. Participants were required to synchronise their press times with those of the virtual partner. The Cued and Interactive blocks were perceptually identical and only differed for the instruction given. **A** – “purple” (Cued instruction) or “same” (Interactive instruction); **B** – “purple” (Cued instruction) or “opposite” (Interactive instruction); **C** – “yellow” (Cued instruction) or “same” (Interactive instruction); **D** – “yellow” (Cued instruction) or “opposite” (Interactive instruction). In 30% of the trials (not shown), the virtual partner changed its initial behaviour 2113 ms (i.e., 66% of the total movement time) after starting its movement and switched, for example, from using its middle finger to stretching the index finger to press the button (Correction trials). In the Interactive condition, but not in the Cued one, Correction trials require participants to adapt their own behaviour to the observed change (i.e., change their own finger) in order to fulfil the request (e.g., to perform a complementary movement), see Video S1.

## Results

### Experimental design and variables

The experimental design of both Experiment 1 and 2 included the within-subjects factors Stimulation (Real, Sham), Block (Interactive, Cued), Trial (Correction, NoCorrection) and Movement (Imitative, Complementary). From the IVR motor interaction task we extracted the following behavioural variables (see STAR Methods for details on the statistical approaches): Asynchrony (i.e., the absolute difference between the participant’s avatar and the virtual partner target-pressing times), Movement Times (i.e., participant’s avatar’s time from start to target pressing), Motor Preparation Times (i.e., participant’s avatar’s reaction time from the “Go” signal to start), First Press Times (i.e., the time at which the subject selected the finger to be used to press the target button for the first time) and Second Press Times (only for Correction trials, i.e., the time at which, during a Correction trial, the subject selected another effector after the first press).

### Statistical approach

The statistical approaches to each variable in Experiment 1 and 2 are detailed in the STAR Methods.

### Experiment 1: Theta tACS reduced interpersonal Asynchrony

The Stimulation (Real, Sham), Block (Interactive, Cued), Trial (Correction, NoCorrection) and Movement (Imitative, Complementary) Aligned Rank Transformation ANOVA showed a significant main effect of Stimulation (*F(*1,19) = 5.99, *p =* .024) (Fig 3, panel a) indicating that interpersonal Asynchrony was reduced during Real, compared to Sham theta stimulation. This main effect was further specified by a Stimulation x Trial x Movement significant interaction (*F(*1,19) = 7.21, *p =* .014) (Fig. 3, panel b). Holm-corrected pairwise comparisons using Wilcoxon signed-rank tests revealed that Asynchrony was reduced in Real theta compared to Sham stimulation in the NoCorrection Complementary condition only (*p =* .022, all other comparisons were not significant *p*s > .088). The analysis also revealed significant main effects of Block (*F(*1,19) = 33.40, *p <* .0001) and Trial (*F(*1,19) = 6.27, *p =* .021), indicating that participants were more synchronous in the Cued, compared to the Interactive block, and in NoCorrection compared to Correction trials.

**Fig. 3.**
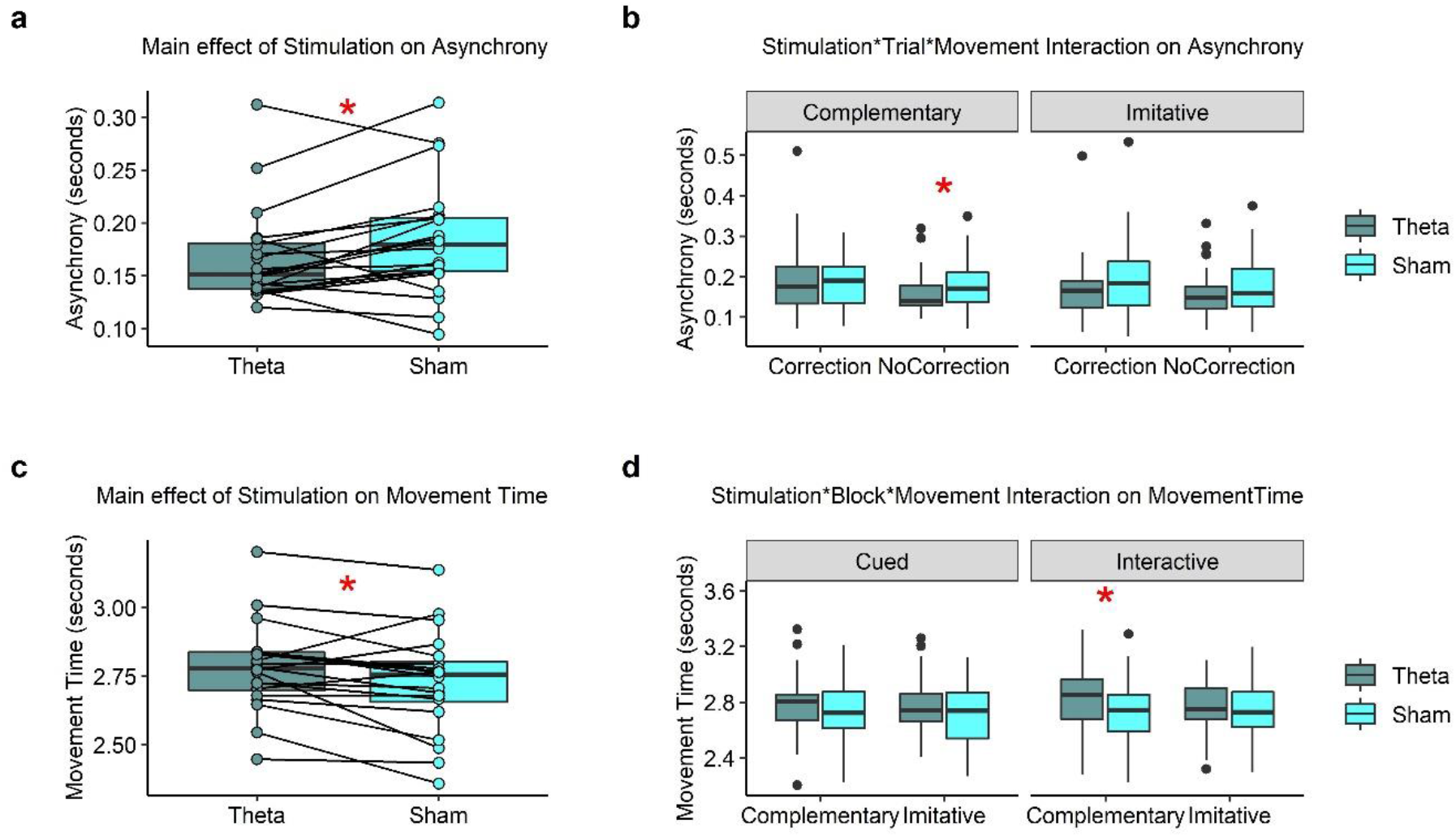
Significant main effect of Stimulation (Panel **a**) and three-level interaction (Panel **b**) on interpersonal Asynchrony. Significant main effect of Stimulation (Panel **c**) and three-level interaction (Panel **d**) on Movement Time. Horizontal lines in the boxes indicate the median, upper and lower borders indicate 1^st^ and 3^rd^ quartile, “whiskers” extend to the farthest points that are not outliers. Colour-filled dots represent individual cases (i.e., participants) in Panels **a** and **c**, black dots represent outliers in Panels **b** and **d**.

### Experiment 1: Theta tACS increased individual Movement Time

The Stimulation (Real, Sham), Block (Interactive, Cued), Trial (Correction, NoCorrection) and Movement (Imitative, Complementary) Repeated Measures Type III ANOVA revealed a significant main effect of Stimulation (*F* (1,19) = 5.76, *p =* .027, η^2^*_p =_* 0.23) indicating that participants’ movement times were increased during Real theta compared to Sham stimulation (Fig. 3, panel c). This main effect was further qualified by the significant Stimulation x Block x Movement interaction (*F* (1,19) = 5.76, *p =* .027, η^2^*_p =_* 0.23). Post hoc tests revealed that theta tACS increased Movement Time with respect to Sham only during Complementary movements in the Interactive block (*p <* .001), while all other comparisons were nonsignificant (all *p*s > .076) (Fig 3, panel d). We also found a main effect of Trial (*F* (1,19) = 16.03, *p <* .001, η^2^*_p =_* 0.45) showing that the movement time increased during Correction compared to NoCorrection trials, and a main effect of Movement (*F* (1,19) = 6.60, *p =* .019, η^2^*_p =_* 0.25) indicating longer movement times in Complementary compared to Imitative trials. There was a significant interaction between Block and Trial (*F* (1,19) = 9.50, *p =* .006, η^2^*_p =_* 0.33). Post hoc tests, however, showed that movement times were longer in Correction, compared to NoCorrection trials both in the Interactive (*p <* .001) and in the Cued block (*p =* .029).

### Experiment 1: Theta tACS reduced Motor Preparation Times in the Interactive block

In the Interactive block, the Stimulation (Real, Sham) and Movement (Imitative, Complementary) Repeated-Measures type III ANOVA revealed a significant main effect of Stimulation (*F* (1,19 = 4.86, *p =* .040, η^2^*_p =_* 0.20) indicating that participants started their movement earlier during Real theta compared to Sham stimulation (Fig. 4, panel a) (all other effects were not significant; effect of Movement (*F* (1,19) = 4.10, *p =* .057, η^2^*_p =_* 0.17)).

**Fig 4.**
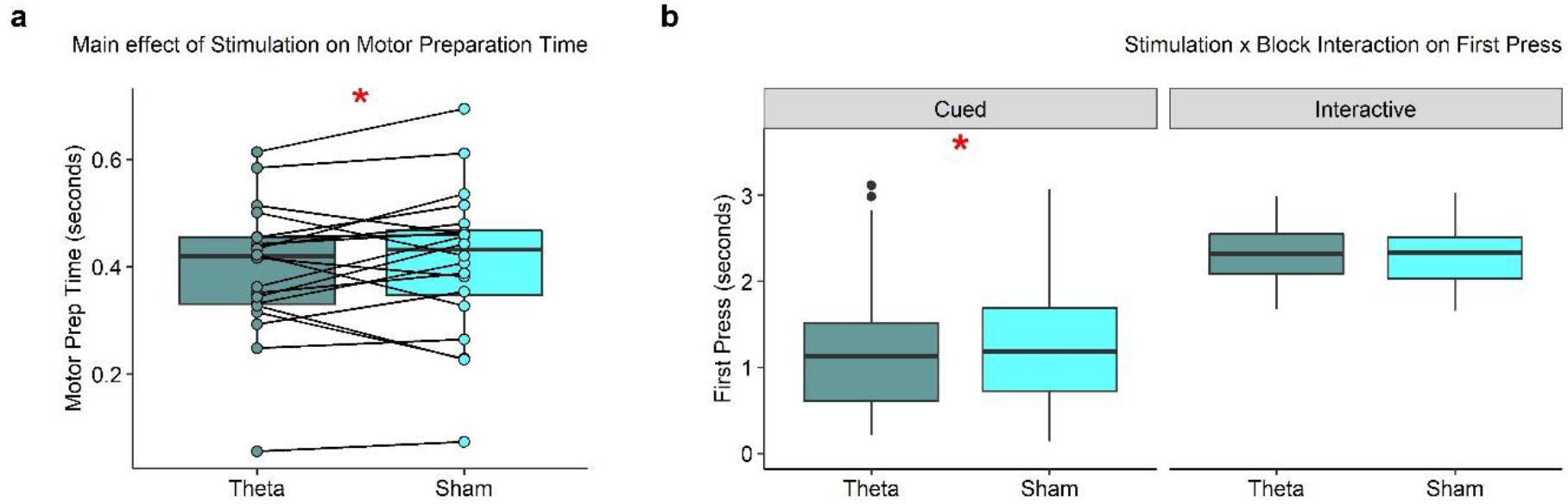
Significant main effect of Stimulation on Motor preparation Time (Panel **a**) and two-level interaction on First Press Time (Panel **b**). Horizontal lines in the boxes indicate the median, upper and lower borders indicate 1^st^ and 3^rd^ quartile, “whiskers” extend to the farthest points that are not outliers. Colour-filled dots represent individual cases (i.e., participants) in Panel **a**, black dots represent outliers in Panels **b**.

Data from the Cued block were analyzed by means of a Wilcoxon signed-rank test between Real and Sham conditions to investigate the effect of Stimulation on Motor Preparation Time. The test was not significant (*p =* .956), indicating that theta tACS did not affect Motor Preparation Time in the Cued block.

### Experiment 1: Theta tACS reduced First Press Times in the Cued block

The Stimulation (Real, Sham), Block (Interactive, Cued), Trial (Correction, NoCorrection) and Movement (Imitative, Complementary) repeated Measures Type III ANOVA revealed significant main effects of Block (*F(*1, 19) = 62.7, *p <* .001, η^2^*_p =_* 0.76), Trial (*F(*1, 19) = 66.9, *p <* .001, η^2^*_p =_* 0.77) and Movement (*F(*1, 19) = 11.06, *p =* .004, η^2^*_p =_* 0.36) indicating that participants selected an effector for the first time earlier in the Cued, compared to the Interactive block, in NoCorrection compared to Correction trials, and in Imitative compared to Complementary trials, respectively. Moreover, a significant Block x Trial interaction revealed that the First Press Time was anticipated in NoCorrection compared to Correction trials in the Interactive block (*p <* .0001) but not in the Cued one (*p =* .835). Finally, a significant Stimulation x Block interaction (*F(*1, 19) = 6.18, *p =* .022, η^2^*_p =_* 0.24) revealed that First Press was anticipated in Real compared to Sham stimulation only in the Cued block (*p =* .001), while no significant difference was observed in the Interactive one (*p =* .45) (Fig. 4, panel b).

### Experiment 1: No effect of theta tACS on Second Press Times in the Interactive block

We analyzed the effect of Stimulation on Second Press Time during Correction trials by means of a paired-sample t-test (two-sided). We found no significant difference between Real and Sham stimulation (t (1,18) = 1.48, *p =* .154), indicating that tACS stimulation did not affect the timing of motor changes during Correction trials.

### Experiment 1: Correlation between behavioural variables

Pearson’s correlations between the behavioural dependent variables were computed with the rcorr.adjust R function, which adjusts the p-value for multiple comparisons using the Holm’s method. Asynchrony and Movement Time were inversely correlated (*r* = −0.62, *p =* .015) (Fig. 5). This indicates that participants who showed longer Movement Time were also better at synchronizing their touch time with the virtual partner. Moreover, we found that Movement Time was inversely correlated with Motor Preparation Time (r = −0.55, *p =* .034). That is, the earlier participants started to move, the longer their movement lasted. Indeed, since Movement Time is computed from the time in which the participant’s Avatar starts to move until when it touches the target, it stands to reason that a reduction in Motor Preparation Time leads to an increase in the movement time. There was no relation between Asynchrony and Motor Preparation Time (r = −0.10, *p =* .999). First Press Time was negatively correlated with Movement Time (r = − 0.60, *p =* .002) and positively correlated with Asynchrony (r = 0.84, *p <* .0001). This indicates that the later was the first effector selection, the shorter was the movement duration and the worst was the synchrony in touching the target with the virtual partner. There was no significant correlation between First Press and Motor Preparation Time (r = − 0.07, *p =* .999).

**Fig 5.**
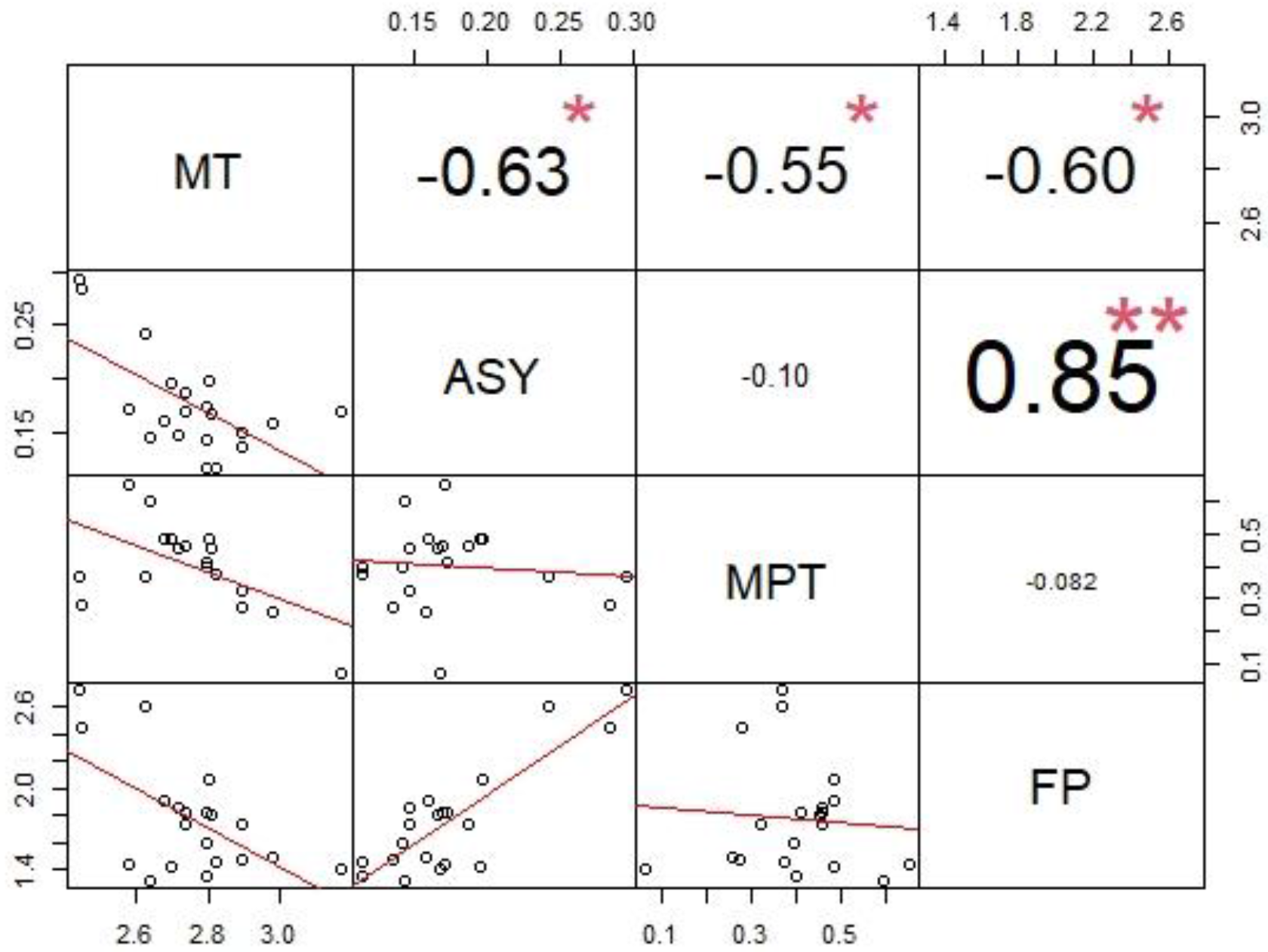
Correlation matrix chart for Experiment 1. On the top of the diagonal line is displayed the value of the correlation coefficient (r) plus the significance level as stars (\*\**p <* 0.01, \**p <* 0.05, adjusted for multiple comparisons with Holm’s method). The font size reflects the magnitude of the correlation coefficient (r). Variable names have been abbreviated: MT = Movement Time, ASY = Asynchrony, MPT = Motor Preparation Time, FP *=* First Press. This plot was generated using the R package “PerformanceAnalytics” (Peterson et al., 2018).

### Experiment 2: Beta tACS did not affect task performance

#### Asynchrony

The Stimulation (Real, Sham), Block (Interactive, Cued), Trial (Correction, NoCorrection) and Movement (Imitative, Complementary) Aligned Rank Transformation ANOVA showed no significant main effect nor interaction (all *F*s < 2.74, all Ps > .072).

#### Movement Time

The Stimulation (Real, Sham), Block (Interactive, Cued), Trial (Correction, NoCorrection) and Movement (Imitative, Complementary) Aligned Rank Transformation ANOVA showed a significant main effect of Trial (*F(*1,19) = 15.53, *p <* .0001) indicating that Movement Times were longer in Correction compared to NoCorrection trials. The Block x Trial (*F(*1,19) = 4.88, *p =* .039) and Block x Movement (*F(*1,19) = 9.09, *p =* .007) interactions resulted to be significant, and were qualified by a Block x Trial x Movement interaction (*F(*1,19) = 4.41, *p =* .049). Wilcoxon signed-rank test revealed that Movement Times were longer in Correction compared to NoCorrection trials only for Imitative trials in the Interactive block (*p <* .001) while the same comparison was not significant for the other conditions (all *ps* > .538). There was no significant difference in Movement Times between Imitative and Complementary trials (all *ps* > .074). Importantly, the main effect of Stimulation was nonsignificant (*F(*1,19) = 0.85, *p =* .366), nor was the Stimulation x Block x Movement (*F(*1,19) = 4.26, *p =* .052) interaction.

#### Motor Preparation Time

In the Interactive block, the Stimulation (Real, Sham) and Movement (Imitative, Complementary) Aligned Rank Transformation ANOVA showed a significant main effect of Movement (*F(*1,19) = 13.81, *p =* .001) indicating that participants started their movement earlier during Imitative compared to Complementary movements. For the Cued block, we used a paired-sample t-test between Real and Sham conditions to investigate the effect of Stimulation on Motor Preparation Time. There was no significant difference between the two (t(1,19) = −1.34, *p =* .195).

#### First Press Time

The Stimulation (Real, Sham), Block (Interactive, Cued), Trial (Correction, NoCorrection) and Movement (Imitative, Complementary) Aligned Rank Transformation ANOVA showed significant main effects of Block (*F(*1,19) = 14.45, *p =* .001), Trial (*F(*1,19) = 21.42, *p <* .001) and Movement (*F(*1,19) = 20.51, *p <* .001). As in Experiment 1, First Press was anticipated in the Cued block, in Imitative movements and in NoCorrection trials. A significant Block x Trial interaction (*F(*1,19) = 11.41, *p =* .003) revealed that First Press was anticipated in NoCorrection compared to Correction trials in the Interactive block (*p <* .0001) but not in the Cued one (*p =* .812). The Stimulation x Block x Movement interaction was nonsignificant (*F(*1,19) = 3.95, *p =* .061).

#### Second Press Time

A paired sample t-test comparing Second Press values in Real and Sham conditions during Correction trials in the Interactive block showed no significant difference between the two stimulation conditions (t(1,13)= −0.74, *p =* .469).

### Experiment 2: Correlation between behavioural variables

Pearson’s correlations between the behavioural dependent variables were computed with the rcorr.adjust R function, which adjusts the p-value for multiple comparisons using the Holm’s method. Results showed the same pattern observed in Experiment 1. Indeed, Asynchrony was negatively correlated to Movement Time (r = − 0.76, *p <* .001). There was no significant correlation between Asynchrony and Motor Preparation Time (r = − 0.19, *p =* .82) nor between Movement Time and Motor Preparation Time (r = − 0.46, *p =* .12). First Press Time was positively correlated to Asynchrony (r = 0.77, *p <* .001) and negatively correlated to Movement Time (r = − 0.80, *p <* .001). There was no significant correlation between First Press Time and Motor Preparation Time (r = 0.13, *p =* .82). See Fig. 6.

**Fig 6.**
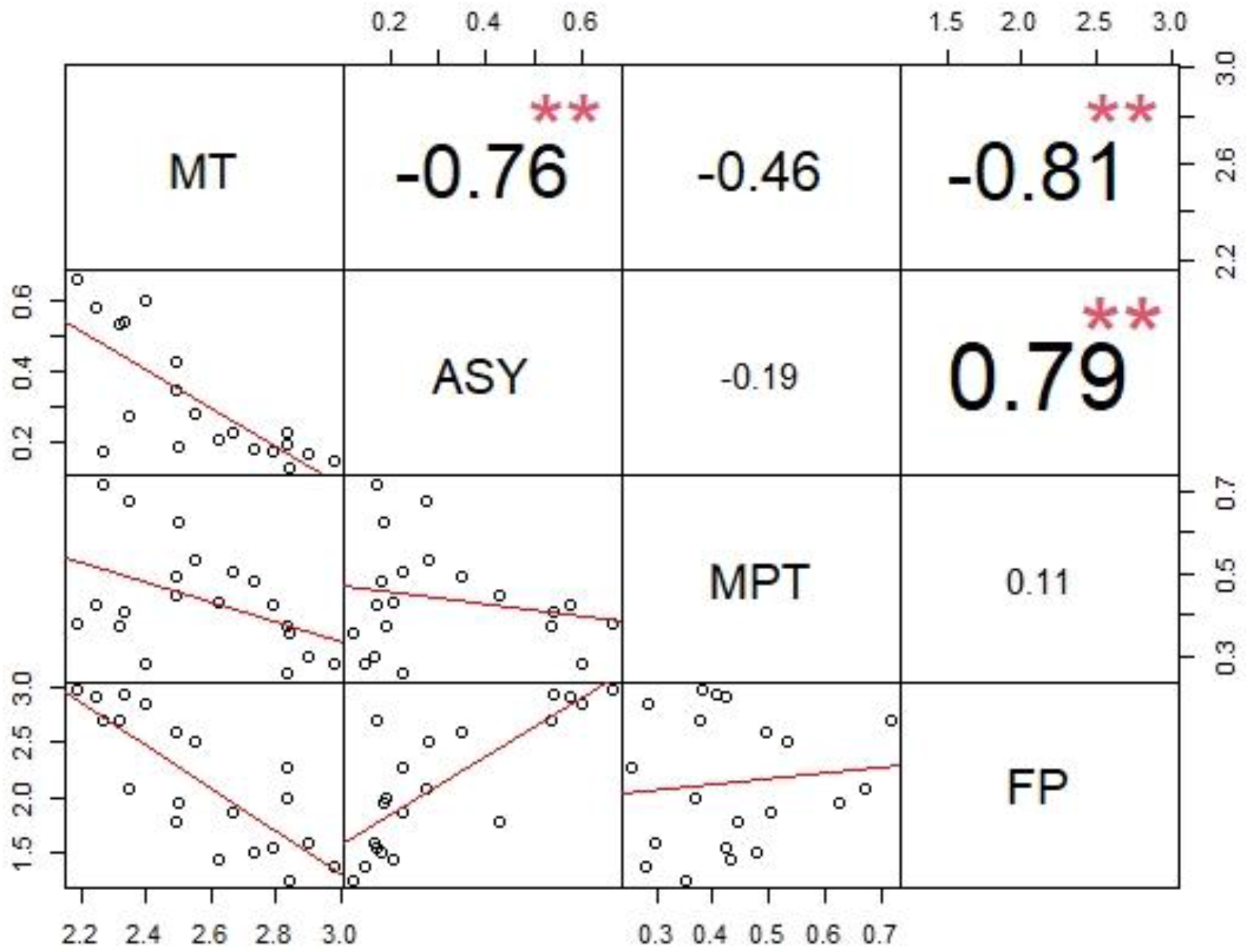
Correlation matrix chart for Experiment 2. On the top of the diagonal line is displayed the value of the correlation coefficient (r) plus the significance level as stars (\*\**p <* 0.01, \**p <* 0.05, adjusted for multiple comparisons with Holm’s method). The font size reflects the magnitude of the correlation coefficient (r). Variable names have been abbreviated: MT = Movement Time, ASY = Asynchrony, MPT = Motor Preparation Time, FP *=* First Press. This plot was generated using the R package “PerformanceAnalytics” (Peterson et al., 2018).

## Discussion

In the present study, we delivered EEG-informed theta (Experiment 1) and beta (Experiment 2) tACS over the frontal midline of participants engaged in a realistic interpersonal motor coordination task with a virtual partner in immersive virtual reality (IVR). In Experiment 1, we tested the hypothesis that previously observed frontal oscillations in the theta range during interpersonal motor interactions (Moreau et al., 2020; 2021) are causally involved in the implementation of motor control during dyadic motor interactions. Experiment 2 served as a control study to ensure that the observed effects were theta-band specific.

We found that frontal theta tACS (Experiment 1) reduced interactors’ Asynchrony (i.e., the absolute difference between participant’s and virtual partner’s target-pressing times, thus resulting in better performance) and increased individuals’ Movement Time (i.e., participant’s time from start to target pressing). These two measures were highly correlated, indicating that participants’ ability to touch the target in synchrony with the virtual partner was correlated with a strategic slowing of the speed at which they were moving their own virtual arm. As detailed in the STAR Methods, the participants’ avatar (i.e., the virtual body observed from a first-person perspective) could move slightly faster than the virtual partner. Thus, being able to reduce the movement speed might have helped participants to reach the target in synchrony with the virtual partner. Although the main effect of theta tACS on Asynchrony was significant (see Fig. 3, panel a), the three-way interaction indicates that the impact of this frequency was stronger during NoCorrection Complementary trials. Similarly, the effect of theta tACS on Movement Time was found to be significant only in the Interactive Complementary condition. Beta frontal tACS, on the other hand, did not affect any behavioural index thus indicating that, in the present experimental set-up, entraining frontal beta oscillation had no effect on the ability to control a virtual agent’s interactive abilities.

### Role of frontal theta for proactive control during interpersonal motor interactions

Although in previous EEG studies (Moreau et al., 2020; 2021), higher theta power was observed after, and locked to, a virtual partner’s correction, increasing frontal theta power in the present study did not have specific effects for Correction trials. It should be noted, however, that rather than being time-locked to the avatar change, in the present study tACS was delivered continuously during the task. This means that, if endogenous midfrontal theta oscillations were present before the time at which the avatar would change its movement, they were most likely entrained by tACS (Helfrich et al., 2014; Witkowski et al., 2016). An increase in frontal theta activity in the pre-trial time window has been reported during several cognitive tasks (Cooper et al., 2015; 2017; 2019; Chang et al., 2017; Adelhöfer et al., 2020) and is correlated to an improvement in task performance in the upcoming trial (Cooper et al., 2019). Pre-trial theta oscillations have been interpreted as the neural correlate of proactive (to be distinguished by reactive) cognitive control, which serves the function of preparing the brain for (un)expected task demands (Stuphorn & Emeric, 2012). In line with an interpretation of theta activity for proactive control it is worth noting that EEG results from Moreau and colleagues’ study (2020) showed an increase in theta power not only in response to an unexpected change in a virtual partner’s movement, but also in No Correction trials when monitoring the partner’s behaviour was essential for successful coordination. Thus, one possible interpretation of the present results is that theta tACS boosted endogenous neural oscillations reflecting proactive cognitive control, which in turn promoted the implementation of a strategic approach by reducing the speed at which participants were moving their avatar. As a result, participants’ ability to “move together” with the virtual partner improved.

### Theta role on early indexes of motor preparation during interpersonal motor interactions

Theta tACS also reduced the time required to initiate the movement (i.e., Motor Preparation Time) in the Interactive block and the time required to select an effector (i.e., First Press Time) in the Cued one. Both indexes reflected participants’ reactivity in the first stages of the trial. Traditionally, midfrontal theta activity has been associated to an increase in reaction times after error commission (Post-Error Slowing, PES, see Cavanagh & Shackman, 2015 for review) that is generally interpreted as the implementation of cognitive control processes to avoid further errors (Notebaert et al., 2009). However, trial-by-trial theta power was also found to be correlated with reduced reaction times in high-conflict trials (Cohen & Donner, 2013). Similarly, studies in which tACS in the theta range was administered to entrain frontal activity reported a beneficial effect on task performance in terms of shorter reaction times (Polania et al., 2012; Lehr et al., 2019; Fusco et al., 2018; 2020). It is generally agreed that the medial prefrontal cortex monitors ongoing behaviour and communicates through long-range slow (e.g., theta) oscillations to other systems, which are in charge of implementing behavioural adaptation (Cavanagh et al. 2012; Cohen 2011; Cohen et al., 2013). In this vein, boosting endogenous theta activity might have reduced reaction times and improved performance through the activation of the performance monitoring system, that is, by increasing the “readiness” of motor areas, supporting the planning of one’s own action based on predictions and monitoring of the other’s action.

Another possible interpretation of our results is that theta tACS helped participants to focus their attention on the motor task and to better control their own movements. Endogenous midfrontal theta activity has indeed been related to sustained attention during cognitive tasks (Sasaki et al., 1996; Onton et al., 2005) and meditation (Aftanas et al., 2001, Tang et al., 2009). However, while this explanation would fit with the general effects of theta tACS that we observed, it fails to account for the condition-specific ones especially since there is no indication that the conditions that benefitted the most from the stimulation were the most difficult ones.

### Limitations of the study

There are potential limitations in this study that should be taken into account in the interpretation of the results. There is still no consensus on the actual duration of tACS aftereffects, which seem highly dependent on the stimulation parameters (Veniero et al., 2015). In particular, the possibility that tACS effects might last after stimulation, affecting plasticity-related neural functioning (Vossen et al., 2015), should be considered. However, the selective effect for theta tACS (compared to beta tACS) in our results suggests that the time between the real and sham stimulation sessions was long enough to abolish real tACS aftereffects and highlight the effects of real compared to sham stimulation. Another potential limitation of our study is the absence of EEG recording after the stimulation, which prevents us to claim that we were, indeed, entraining midfrontal theta power, the only indication of such effect being the behavioural effects found after theta, and not beta, tACS.

### Conclusion and future directions

Recent studies have generated interest in the role of midfrontal oscillations in goal-directed interpersonal motor interactions (Ninomiya et al., 2018; 2020) and a specific role of humans’ frontal theta during interpersonal reaching interactions (Moreau et al., 2020). In the present study we investigated this issue from a causative approach, capitalizing on the combination of tACS and virtual reality. We found that frontal theta (but not beta) tACS improved participants’ ability to synchronize their movements with those of a virtual partner, possibly by hyper-activating the performance monitoring system. Our results hint to a beneficial effect of theta tACS on motor behaviour in a social context and could pave the way toward non-invasive brain stimulation approaches used in interactive situations in treating motor and cognitive disorders (Candidi et al., 2017; Pezzetta et al., 2021). Moreover, to the best of our knowledge this is the first study in which tACS was administered when participants are immersed in a virtual reality scenario. Thus, our research shows the feasibility of brain stimulation-IVR combination and can be considered as a first step in this direction. Future research should consider the potential benefits of EEG recording following theta tACS stimulation to measure its physiological effects and their relationship with behaviour.

## Author Contributions

SB, VE, QM, MC designed the experiment; GT realized the virtual scenario and programmed the task; SB and DGO recorded the data; SB, DGO and QM analyzed the data; SB, DGO, VE, QM and MC wrote the manuscript; MC raised the funding.

## Acknowledgments

MC was funded by the Italian Ministry of Health (Ricerca Finalizzata, Giovani Ricercatori 2016, protocol number GR-2016-02361008), Sapienza University (Progetti di Ricerca Grandi 2020). VE was supported by the Fondazione Umberto Veronesi.

---

**Supplementary Video S1:** https://youtu.be/jzdU6FyXrUQ

## Supplemental figure titles and legends

**Table S1**– Embodiment Questionnaire. Items Q1, Q2, Q4 and Q5 measure Ownership (e-O; i.e. the feeling that the virtual body belong to the observer) and Agency (e-A; the feeling to be in control of the virtual body), respectively. Items Q3 and Q6 were control items for FO and A. Items Q1 and Q2 were averaged together to create the Ownership item. Items Q4 and Q5 were averaged to create the Agency item.

**Figure S1** - Modelling of electric fields using Realistic vOlumetric-Approach to Simulate Transcranial Electric Stimulation (ROAST, Huang et al., 2019).

**Fig. S2**– Experiment 1 (theta tACS) - Histogram plot describing the number of reported cases for each level of the five-points scale in the tACS- induced sensations questionnaire after Real and Sham stimulation.

**Fig. S3** – Experiment 2 (beta tACS)- Ratings for the Agency item were significantly different in Real compared to Sham condition (*p =* .022), indicating that participants reported lower feeling of Agency in Real compared to Sham tACS blocks.

**Fig. S4 –** Experiment 2 (beta tACS) - Histogram plot describing the number of reported cases for each level of the five-points scale in the tACS- induced sensations questionnaire after Real and Sham stimulation.

## STAR Methods

### Resource availability

#### Lead contact

Further information and requests for resources should be directed to and will be fulfilled by the lead contact Sarah Boukarras

#### Materials availability

This study did not generate new unique reagents.

#### Data and code availability

The datasets and codes generated during this study are available at OSF repository https://osf.io/c6sb3/?view_only=6493544897c3480990a8a5e58b05d11b.

### Experimental model and subject details

Details on the subjects are reported in the Participants section of the STAR Methods.

### Method details

#### Participants

Twenty-one healthy participants were recruited in Experiment 1 and twenty-two in Experiment 2. One subject was excluded from Experiment 1 for reporting excessive discomfort induced by tACS. Two subjects were excluded from Experiment 2 because they reported motion sickness induced by Immersive Virtual Reality (1 subject) and anomalies in the resting-state EEG Alpha peak shape (1 subject), respectively. The final sample for Experiment 1 (theta tACS) included twenty (20) right-handed participants (mean age = 25.4 years; range = 21 – 36; 10 M, 10 F). The final sample for Experiment 2 (beta tACS) also included twenty (20) right-handed participants (mean age = 23.3 years, range 20 – 33; 12 M, 8 F). The sample size was determined from previous studies that employed tACS with a similar design (Onoda et al., 2017; Zaehle et al., 2010). All participants gave written informed consent to participate in the study. Exclusion criteria were any declared neurological or psychiatric issues. Suitability to receive non-invasive brain stimulation was assessed by a standardized questionnaire (Antal et al., 2017). All participants had normal or corrected-to-normal vision and were naïve as to the purpose of the experiment. The experimental procedure was approved by the Ethics Committee of IRCCS Santa Lucia Foundation of Rome and was performed in accordance with the ethical standards of the 1963 Declaration of Helsinki.

#### Experimental stimuli

The virtual environment and avatars were designed by means of 3DS Max 2017 (Autodesk, Inc.) and IClone 7 (Reallusion, Inc.) respectively, and implemented in Unity 5 game engine software (http://unity.com). The virtual environment was presented by means of the Oculus Rift Head Mounted Display (HMD; www.oculus.com) having 110° field-of-view (diagonal FOV) with a resolution of 2160 x 1200. The virtual scene consisted of a real-size room (1:1 scale), two virtual avatars, one seen in first person-perspective, 1PP (i.e., participant’s avatar), one in third person perspective (i.e., partner’s avatar) sitting on opposite sides of a table and a virtual grey panel placed between the avatars so that the faces were not visible (similar to the VR scenario used in Fusaro et al., 2019 and Moreau et al 2021). In front of both avatars, at the center of the table, there were two virtual buttons, colored purple and yellow, and a virtual LED light that could either turn red or green. Participants observed their virtual body and the scenario from 1PP through HMD and used the right Oculus Touch controller to control the movement of the right arm of their virtual body in real time. In particular, participants could i) move their virtual hand forward in space and control its velocity by using the analogue stick of the Oculus touch controller with right thumb and ii) animate the right index or middle finger by pressing the Oculus touch controller’s up and down trigger button with their own right index and middle finger, respectively. During the experiment, the virtual scenario was rendered in both HMD and a computer screen, such that the experimenter could observe and assist the participants.

#### Immersive Virtual Reality Motor Interaction task

The IVR Motor Interaction Task (Figure 3) comprised two blocks: Interactive and Cued (see Video S1) that differed for (i) the instruction received and (ii) the type of interaction required. In particular, in the *Interactive* block, participants were asked to reach and press one of the two virtual buttons as synchronous as possible with the virtual partner while performing either an imitative (instruction “same”) or a complementary (instruction “opposite”) movement with respect to the virtual partner’s (e.g., if the instruction, received before each trial, was ‘opposite’ and the virtual partner raises the index finger to press the purple button, then participant needed to raise the middle finger to press the yellow button). In the *Cued* block, participants had to synchronize their reach-to-press movements with those of the virtual partner as in the Interactive block but were in this case instructed to press either the “purple” (with the index finger) or “yellow” (with the middle finger) buttons, regardless of which action the avatar was performing. Thus, in the Interactive block participants needed to predict and monitor the action of the virtual partner in order to perform their own action, while action prediction and monitoring were not needed during the Cued condition, where participants already knew what action to perform. Importantly, in 30% of the trials of both Interactive and Cued blocks, the virtual partner changed its initial behaviour 2113 ms after starting its movement (i.e., 66% of the total movement time), namely it switched from using its middle finger to stretching the index finger to press the button (Correction trials). It is important to note that in the Interactive condition, but not in the Cued one, Correction trials require participants to adapt their own behaviour to the observed change (i.e., change their own finger) in order to fulfil the request (e.g., to perform a complementary movement). However, the presence of Correction trials in the task boosted the need for participants to monitor the virtual partner’s action in all the trials, since participants did not know in advance whether a specific trial would be a Correction trial. Each trial started with an acoustic ‘go’ signal (“beep”) delivered through the HMD’s headphones. Both avatars started with their hands closed and placed in the center of table’s midline. After the go signal was delivered, the participant and the virtual partner started moving (virtual-partner total movement time lasting 3170 ms). The virtual partner’s starting time was 40 ms after the “go” signal, then 1056 ms after it started moving (i.e., 33% of the whole movement time), the virtual partner would raise a finger in order to press the associated button (index finger for purple button and middle finger for yellow button, see Fig 3, right panel). It was the participant’s duty to control his/her avatar’s right hand with the Oculus Touch controller to reach and press one of the two buttons as synchronously as possible with the virtual partner. With the analogue stick of the controller, participants could move their avatar’s arm forward and regulate its velocity (i.e., velocity was proportional to the force applied by their thumb) and, by pressing the index and middle trigger button of the controller, they could raise either the virtual index or the virtual middle finger of their avatar. Depending on the Asynchrony (i.e., absolute time difference between the two pressing times) the virtual LED light could turn either green (‘win’ trial) or red (‘fail’ trial). A staircase procedure was adopted to make the task more challenging for pressing the button synchronously with the virtual partner: after each ‘win’ trial the minimum time difference to turn the light green was reduced of 50 ms (e.g., from 200 ms to 150 ms), while in the case of ‘fail’ trials, the time window was increased of 50 ms (e.g., from 200 to 250 ms). The trial ended 2 seconds after the LED visual feedback. The avatar’s total movement time (i.e., the time from start to touch) lasted approximately 3.2 seconds. Each of the two tasks (i.e., Cued and Interactive) comprised 68 trials, of which 20 were Corrections (10 Opposite, 10 Same) and 48 were NoCorrection (24 Opposite, 24 Same).

#### Procedure

The study comprised two different phases: 1) the extraction of the individual EEG frequency at rest and 2) the execution of the IVR task during tACS stimulation (i.e., experimental session, see Fig 1). During Phase 1, at their arrival at the laboratory, participants went through an EEG resting-state recording session (see details below) where they were asked to sit in a quiet room and stay still with their eyes closed for 5 min. Then, participants had a small break (∼ 20 minutes), during which the individual-frequency information was extracted. Prior to the tACS session, subjects’ scalp was measured to determine FCz and Pz positions according to the International 10-10 EEG layout. The areas of interest were cleaned with a cotton swab soaked in ethyl alcohol in order to reduce the skin’s conductance and marked with a marker. The two tACS electrodes were then fitted through an EEG-cap over the head of the participants, with the side toward the skin coated with electro-conductive gel. During Phase 2, participants first had the opportunity to familiarize with the tACS-induced physical sensations (i.e., itching, heat) by receiving 15 seconds (5 s ramp-up, 5 s stimulation, 5 s ramp-down) of tACS at 13 Hz and were asked to report any physical sensation or discomfort. If no discomfort was reported, participants entered in the IVR scenario by wearing the HMD where they would observe a virtual body from a first-person perspective (1PP) and the virtual partner from a third-person perspective (3PP) point of view (see Experimental stimuli to detailed description of IVR scenario and stimuli).

Before starting the experiment, participants underwent the IVR-Calibration, -Familiarization and -Training phases, respectively. In the IVR-Calibration phase, the perspective point-of-view of each participant was adjusted to match the virtual body, observed in 1PP, with individual positioning in order to obtain the best spatial-match between the participant’s real and virtual body. In the IVR-Familiarization phase, participants were invited to look both at their virtual body (the participant’s avatar) and at the environment, and to verbally describe what they were seeing (∼30 sec) (Tieri et al. 2015). During the IVR-Training phase, which was provided at the beginning of each of the experiment, participants completed 10 trials of the IVR Motor Interaction Task. The experimental phase of both Experiment 1 and Experiment 2 consisted of two Stimulation blocks: 1) real tACS stimulation (at individual’s theta in Experiment 1, or beta in Experiment 2) and 2) sham tACS block, presented in counterbalanced order, each of which comprised two Task blocks (Interactive and Cued, order counterbalanced). At the end of each Stimulation block there was a 5-minute break during which participants completed the Embodiment Questionnaire and a standardized questionnaire measuring tES-induced physical sensations (Fertonani et al., 2015).

#### Embodiment questionnaire

After each experimental block, a black panel with a horizontal green line (VAS scale, 60 cm length, left and right extremity marked as “0” and “100” respectively) was presented in the virtual scenario. In order to assess the degree to which participants experienced the illusory Feeling of Ownership (FO), i.e., the feeling that the virtual body belong to them, and Agency (A), i.e., the feeling to be in control of the virtual body’s action, a 6-item questionnaire adapted from previous studies (Botvinick & Cohen, 1998; Tieri et al. 2015; 2017; Villa et al., 2018; 2020; Provenzano et al., 2019; Fusco et al. 2020b; Fusaro et al., 2021) (Table 2) was used. The questionnaire included three items concerning the FO (Q1–2 experimental, Q3 control) and Agency (Q4–5 experimental, Q6 control), respectively. Participants were asked to move a vertical bar along the horizontal VAS line by using the analogue stick of the right Oculus touch controller in order to answer the items reported in Table S1.

Experimental items of the Embodiment Questionnaire (Q1 and Q2 for Ownership, Q4 and Q5 for Agency) were averaged together to create two Embodiment ratings (Agency and Ownership) to be compared against the Control items (Agency_Control and Ownership_Control). Since the questionnaire was administered at the end of each stimulation session (i.e., after an Interactive + Cued block), we investigated the effect of Real and Sham stimulation on Embodiment ratings and Control items. For each measure, ratings of Agency, Ownership and their respective Control items were analysed through a non-parametric Friedman ANOVA and follow-up comparisons with Wilcoxon paired tests, Bonferroni corrected for multiple comparisons.

#### tACS-induced sensations questionnaire

After each experimental session, participants were asked to fill a questionnaire designed to measure the presence and intensity of physical sensations induced by electrical stimulation (Fertonani et al., 2015). Participants rated the intensity of each sensation (Itching, Pain, Burning, Warmth/Heat, Pinching, Metallic/Iron Taste, Fatigue) on a 5-points scale from None (0) to Strong (5). At the end of the experiment, participants were asked whether they believed to have received a placebo stimulation in the first or in the second stimulation session. Items from the Transcranial Electrical Stimulation-induced (TES-induced) sensations questionnaire (Itching, Pain, Burning, Heat, Pinging and Fatigue) were analysed with paired-sample Wilcoxon signed rank tests to check for within-subjects differences between Real and Sham stimulation conditions. Data from the Iron Taste item were not analysed, as only three responses in the whole dataset were higher than zero.

Statistical analyses and results for the Embodiment Questionnaire and the tES-induced sensations questionnaire are reported in the Supplementary Materials.

#### EEG protocol

EEG signals were recorded via Neuroscan SynAmps RT amplifier system, from an elastic headband (Electro-Cap International) EEG arranged according to the International 10-10 EEG System with 58 scalp electrodes (Compumedics, ltd). EEG was recorded using following channels: Fp1, Fpz, Fp2, AF3, AF4, F7, F5, F3, F1, Fz, F2, F4, F6, F8, FC5, FC3, FC1, FCz, FC2, FC4, FC6, T7, C5, C3, C1, Cz, C2, C4, C6, T8, TP7, CP5, CP3, CP1, CPz, CP2, CP4, CP6, TP8, P7, P5, P3, P1, Pz, P2, P4, P6, P8, PO7, PO3, PO1, POz, PO2, PO4, PO8, O1, Oz, O2. The amplifier hardware band-pass filter was 0.01–200 Hz and the sampling rate was 1000 Hz. Impedances were lowered below 5 kΩ using electro-gel. Reference electrodes were applied to the left (digital reference) and right (physical reference) earlobes, and all electrodes were re-referenced offline to the average of both.

#### tACS protocol

Electrical stimulation was delivered via two circular sponge-based rubber electrodes (Sponstim, 25 cm, Neuroelectrics, Barcelona, Spain) soaked in saline water (NaCl) and connected to a rechargeable battery-operated stimulator system (Starstim/Enobio, Neuroelectrics, Barcelona, Spain) which in turn was controlled via Bluetooth by a dedicated software (Neuroelectrics Instrument Controller – NIC, Neuroelectrics, Barcelona, Spain). Electrodes were placed over the midline at FCz and Pz (International 10-20 System) beneath an EEG cap, see. We modelled the electric field strength for our montage using the “Realistic vOlumetric-Approach to Simulate Transcranial Electric Stimulation” (ROAST), (Huang et al., 2019), see Fig S1. Finite-element modelling was performed using the MNI152 averaged head. Participants received individualized theta (Experiment 1, mean Hz = 5.5 ± 0.65) or beta (Experiment 2, mean Hz = 17.6 ± 2.5) tACS with an intensity of 2 mA (peak-to-peak) while engaged in the IVR-based Motor Interaction task. Impedance was kept below 5KΩ. Stimulation/Task blocks lasted approximately from 9’ to 9’30’’. During each block, the current was ramped up for 5 seconds before starting the task and ramped down for 5 seconds after the task was completed. In half of the blocks, participants received sham stimulation which included 5 seconds of ramp up, 20 seconds of AC and 5 seconds of ramp down.

#### Resting state EEG data analysis

The EEG data analysis was performed using the FieldTrip toolbox for EEG/MEG (Oostenveld et al, 2011; Donders Institute for Brain, Cognition and Behaviour, Radboud University, the Netherlands. See http://fieldtriptoolbox.org). In order to extract the individual peak frequencies, the five-minute resting state recordings were segmented into epochs of 4 seconds (Pahor & Jaušovec 2014). Independent Component Analysis (Jung et al., 2000) was computed to identify and remove eye movements and muscular artifacts. An average of 0.94 components (SD = 0.75) per subject were removed and 69.53 artifact-free epochs (SD = 4.17) per participant was kept. Data were band-pass filtered at 1-70Hz and a Fast Fourier Transformation with 0.25 Hz resolution was performed to derive estimates of absolute spectral power (Pahor & Jaušovec 2014). We first identified the individual Alpha peak frequency (IAF) (*M*_IAF_ = 10.64, *SD*_IAF_ = 0.66). Following Methods from Klimesch (1999), individual Theta frequency (ITF, Experiment 1) was extracted by choosing the highest peak between IAF 4.0 and 6.0 Hz range (*M*_ITF_ = 5.5, *SD*_ITF_ = .65). For individual Beta frequency (IBF, Experiment 2), the peak between 12.5 Hz and 22.5 Hz was chosen (*M*_ITF_ = 17.5, *SD*_ITF_ = 2.54). The calculated peaks were rounded-up to 0.5 Hz and were visually inspected and confirmed (Klimesch 1999; van Driel et al., 2015). For 7 participants, the automatic calculation placed the ITF to lower frequencies outside the Theta range due to the presence of pink noise, in such cases the peak frequency was manually determined as to the largest frequency within the Theta range.

### Quantification and statistical analysis

As a first step, for each behavioural variable (Asynchrony, Movement Times, Motor Preparation Times, First and Second Press Times) we removed trials in which participants i) failed to follow the instructions (i.e., Same or Opposite for the Interactive Block and Purple or Yellow for the Cued), ii) failed to touch the target. Moreover, from Second Press Times we removed Correction trials in which the participant did not perform a motor change (i.e., he/she selected an effector only once thus not switching from one finger to the other) and, only from First Press, we removed NoCorrection trials in which the participant performed a correction (i.e., he/she selected both effectors).

From these new datasets (Experiment 1 and 2), we removed outliers following the Tukey’s interquartile range (IQR) approach (https://goo.gl/4mthoF). We used the Shapiro-Wilk test from the R package “rstatix” (Kassambara, 2020) to check if our variables were normally distributed. When the assumption of normality was not met, we conducted non-parametric repeated-measures factorial analysis of variance on the aligned and rank-transformed data with the R package ARTool (Kay & Wobbrock, 2016). The Aligned Rank Transform (ART) procedure overcomes the limitation of the more common (one-way) Friedman ANOVA by allowing the examination of interaction effects and is thus more suitable for repeated-measures designs with more than one within-subjects factors (Wobbrock et al., 2011). Post hoc tests were carried using non-parametric Wilcoxon signed-rank tests corrected for multiple comparisons using Holm’s sequential Bonferroni procedure (Holm 1979). If the data were normally distributed, we conducted repeated-measures Type III ANOVA tests with the *anova_test* function from *rstatix* (Kassambrara, 2020) package. Normality of model residuals was visually inspected using Q-Q plots and histograms.

For Asynchrony, Movement Time and First Press, the within-subjects effects included Stimulation (Real, Sham), Block (Interactive, Cued), Trial (Correction, NoCorrection) and Movement (Imitative, Complementary).

Motor Preparation Times were analysed separately for the Interactive and Cued block, since in the two conditions some of the factors did not have any meaning. Indeed, in the Cued block the instruction was to press either the Purple or the Yellow target and participants could not know whether they were to perform a Complementary or an Imitative trial at movement onset. Within-subjects effects thus included Stimulation and Movement for the Interactive model and Stimulation for the Cued one. Furthermore, the effect of Trial was not included in any of the two analyses since at the beginning of the trial participants could not know whether the virtual partner would correct its movement.

Data from Second Press Time were analysed through a paired-sample t-test between Real and Sham conditions.

To investigate the relation between the different variables, we ran Pearson correlations between Asynchrony, Movement Time, Motor Preparation Time and First Press Time averaged across conditions. The p-values were corrected for multiple inference using Holm’s method.

### Supplemental item titles

Video S1

