## Supplementary Materials for "Midfrontal theta tACS facilitates motor coordination in dyadic human-avatar motor interactions"

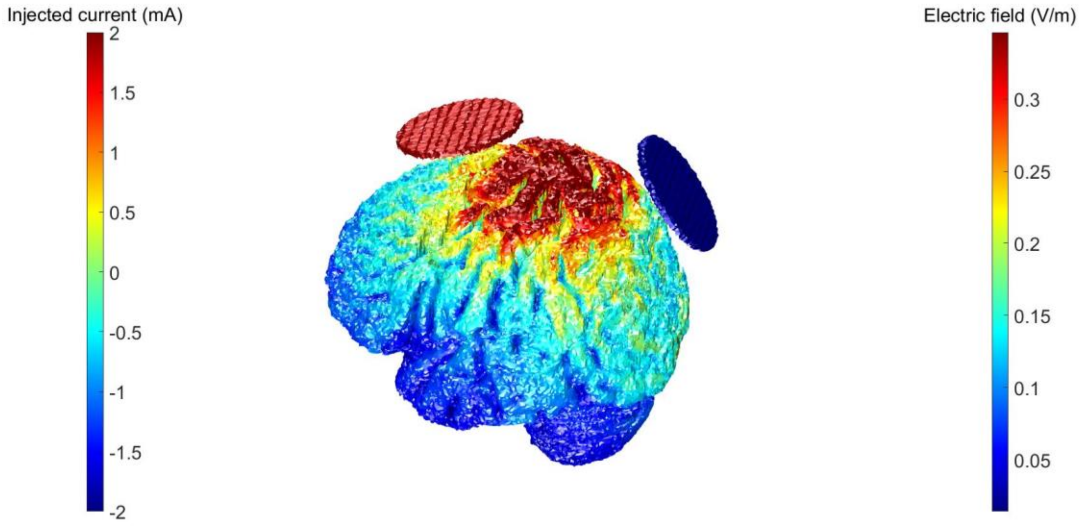


**Figure S1** - Modelling of electric fields using Realistic Volumetric-Approach to Simulate Transcranial Electric Stimulation (ROAST, Huang et al., 2019).

| **Index** | **Item** | | |
|---|---|---|---|
| *e-O* | **Q1** | I felt as if I were looking at my own hand | |
| *e-O* | **Q2** | I felt as if the Virtual Hand were my hand | Ownership |
| *c-O* | **Q3** | It felt as if I had more than one right hand | Ownership Control |
| *e-A* | **Q4** | It felt as if the movements of the Virtual Hand were my own movements | |
| *e-A* | **Q5** | I felt as if I could have caused a/the movement of the Virtual Hand | Agency |
| *c-A* | **Q6** | I felt as if the Virtual Hand were controlling me | Agency Control |

Table S1– Embodiment Questionnaire. Items Q1, Q2, Q4 and Q5 measure Ownership (e-O; i.e. the feeling that the virtual body belong to the observer) and Agency (e-A; the feeling to be in control of the virtual body), respectively. Items Q3 and Q6 were control items for FO and A. Items Q1 and Q2 were averaged together to create the Agency item. Items Q4 and Q5 were averaged to create the Ownership item.

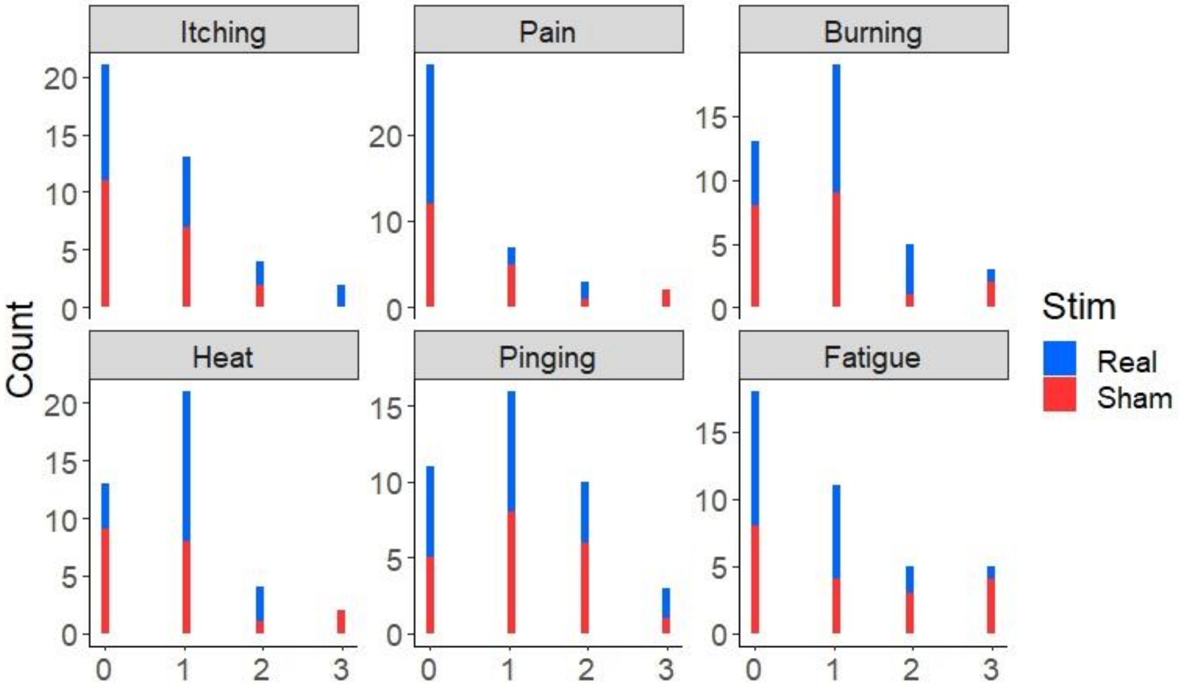


Fig. S2– Experiment 1 (theta tACS) - Histogram plot describing the number of reported cases for each level of the five-points scale in the tACS- induced sensations questionnaire after Real and Sham stimulation.

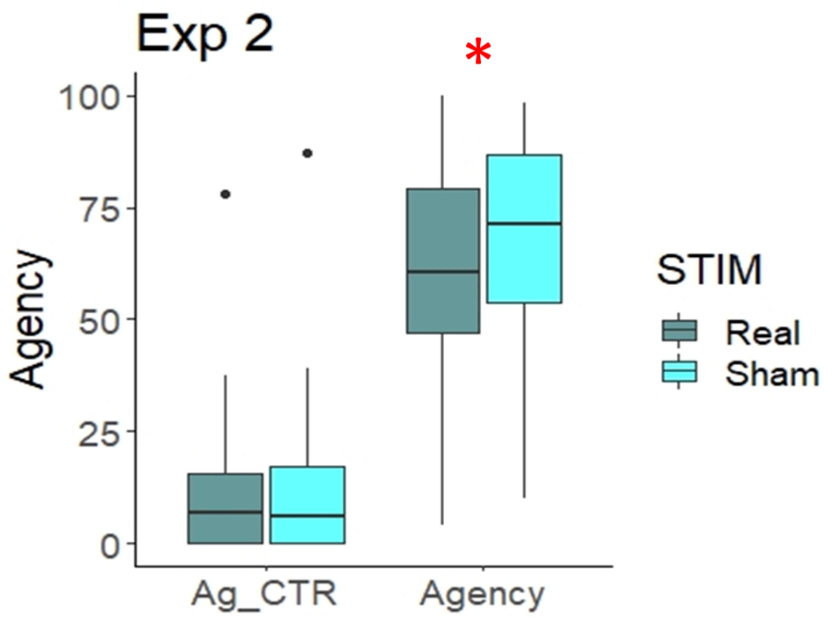


Fig. S3 – Experiment 2 (beta tACS)- Ratings for the Agency item were significantly different in Real compared to Sham condition ($p$ = .022), indicating that participants reported lower feeling of Agency in Real compared to Sham tACS blocks.

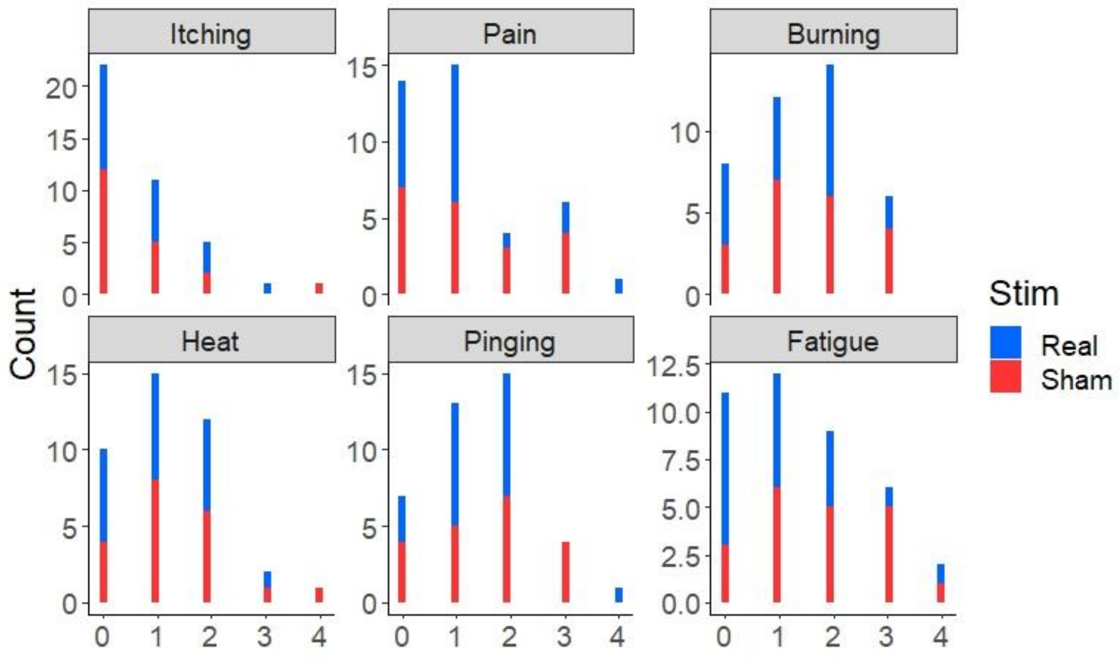

Fig. S4 – Experiment 2 (beta tACS) - Histogram plot describing the number of reported cases for each level of the five-points scale in the tACS- induced sensations questionnaire after Real and Sham stimulation.

### Supplementary Results

*Experiment 1 - Embodiment Questionnaire*

*Ownership.* Friedman ANOVA on Ownership ratings was significant ($\chi^2$(N=19, df = 3) = 15.16, $p$ = .001). Wilcoxon tests showed that ratings for the Ownership items were significantly higher than ratings for the Ownership_Control items both in the Real ($p$ = .001) and Sham ($p$ = .004) conditions. Ratings for Ownership and Ownership_Control items were not different in Real compared to Sham stimulation (all *Ps* > .605), indicating that tACS had no effect on the feeling of Ownership.

*Agency.* Friedman ANOVA on Agency ratings was significant (χ2(N=19, df = 3) = 39.74, $p$ < .0001). Wilcoxon tests showed that ratings for the Agency items were significantly higher than ratings for the Agency_Control items both in the Real ($p$ < .001) and in the Sham ($p$ < .001) conditions. Ratings for Agency and Agency_Control items were not different in Real compared to Sham condition for experimental nor for control items (all *Ps* > .593), indicating that tACS had no effect on the feeling of Agency.

*Experiment 1 – tACS-induced sensations questionnaire.*

Paired-sample Wilcoxon signed rank tests failed to show any significant difference between Real and Sham (all ps > .240), indicating that participants did not report stronger sensations

(Itching, Pain, Burning, Heat, Pinging and Fatigue) in Real compared to Sham stimulation (see Fig. S2).

*Experiment 2 - Embodiment Questionnaire*

*Ownership.* Friedman ANOVA on Ownership ratings was significant ($\chi 2(N=19, df = 3) = 12.63$, $p = .005$). Wilcoxon tests showed that ratings for the Ownership items were significantly higher than ratings for the Ownership_Control items both in the Real ($p = .016$) and in the Sham ($p < .001$) conditions. Ratings for Ownership and Ownership_Control items were not different in Real compared to Sham condition (all *Ps* > .078), indicating that tACS had no effect on the feeling of Ownership.

*Agency.* Friedman ANOVA on Agency ratings was significant ($\chi 2(N=19, df = 3) = 41.62$, $p < .0001$). Wilcoxon tests showed that ratings for the Agency items were significantly higher than ratings for the Agency_Control items both in the Real ($p < .001$) and in the Sham ($p < .001$) conditions. Ratings for the Agency items were significantly different in Real compared to Sham condition ($p = .022$), indicating that participants reported lower feeling of Agency in Real compared to Sham tACS blocks (see Fig. S3). Conversely, there was no significant difference between Real and Sham for the Agency_Control item ($p = .826$).

*Experiment 2 - tACS-induced sensation questionnaire*.

Paired-sample Wilcoxon signed rank tests revealed a significant difference between Real and Sham stimulation for Fatigue rating ($p = .033$). All other comparisons were nonsignificant (all *p*s > .305). See Fig S4.

**Supplementary Discussion**

In accordance with previous research (Tieri et al. 2015; 2017; Villa et al., 2018), ratings for both Agency and Ownership were significantly higher than ratings given to Control items, indicating that participants were experiencing feelings commonly associated to the embodiment of a virtual avatar. Specifically, they felt that the virtual arm belonged to their body (Ownership) and that they could control the movement of the virtual arm (Agency). This indicates that the task might be effective in recreating a real-life motor interaction scenario in Virtual Reality. In Experiment 2, we observed that participants' ratings of Agency were reduced in Real (beta) tACS compared to Sham stimulation. A recent study combining fMRI and TMS have identified a brain network including the pre-supplementary motor area (SMA) and the dorsal parietal cortex associated to the sense of agency (Zapparoli et al., 2020). By delivering high-frequency (10 Hz) TMS over the pre-SMA, the researchers were able to disrupt the feeling of agency during goal-directed actions as measured by intentional binding. Since in our study tACS was delivered over the FCz EEG position, it is possible that the electrical current flow reached the SMA (electrodes in tDCS/tACS studies targeting SMA are placed anterior to Cz, see for example Carlsen et al., 2015) and interfered with the feeling of agency. The fact that this reduction was only visible after beta tACS is somehow surprising as it suggests an opposite role of fronto-parietal beta power in supporting the feeling of ownership compared to the results of an EEG study that showed that the induced belief of agency during a motor task was associated to increased beta-mediated connectivity between motor and parietal areas (Buchols et al., 2019). Further research is needed in order to clarify the role of distinct neural oscillations in the sense of agency. Another possibility to consider is that in our sample the reduction in the sense of agency was caused by an increase in fatigue levels triggered by beta tACS (see paragraph *Experiment 2 - tACS-induced sensation questionnaire*).
